# Reward influences the allocation but not the availability of resources in visual working memory

**DOI:** 10.1101/2021.06.08.447414

**Authors:** James A. Brissenden, Tyler J. Adkins, Yu Ting Hsu, Taraz G. Lee

## Abstract

Visual working memory possesses strict capacity constraints which place limits on the availability of resources for encoding and maintaining information. Studies have shown that prospective rewards improve performance on visual working memory tasks, but it remains unclear whether rewards increase total resource availability or rather influence the allocation of resources without affecting availability. Participants performed a continuous report visual working memory task with oriented grating stimuli. On each trial, participants were presented with a priority cue, which signaled the item most likely to be probed, and a reward cue, which signaled the magnitude of a performance-contingent reward. We showed that rewards decreased recall error for cued items and increased recall error for non-cued items. We further demonstrated that rewards produced a tradeoff in the probability of successfully encoding a cued versus a non-cued item rather than a tradeoff in recall precision or the probability of binding errors. Lastly, we showed that rewards only affected resource allocation when participants were given the opportunity to engage proactive control prior to encoding. These findings indicate that rewards influence the flexible allocation of resources during selection and encoding in visual working memory, but do not augment total capacity.

## Introduction

Visual working memory is a goal-directed process that is severely capacity limited. Capacity constraints necessitate the ability to select or prioritize a subset of available information (i.e., the flexible allocation of resources). While it is well established that resources can be flexibly allocated across items held in working memory (van den Berg et al., 2012; Fougnie et al., 2012; van den Berg and Ma, 2018), it is currently unclear whether there is also flexibility in the total amount of resources that can be allocated at different points in time. It has been shown that monetary rewards facilitate a broad range of processes. The prospect of receiving reward could potentially drive flexibility in the allocation of visual working memory resources across time.

Monetary incentives increase both the speed and accuracy of movements, seemingly violating the speed-accuracy tradeoff predicted by optimal control theory (Takikawa et al., 2002; Manohar et al., 2015, 2018; Anderson et al., 2020; Adkins et al., 2021). Participants also benefit from rewards in the performance of cognitive control tasks, exhibiting both reduced reaction time and increased accuracy (Hübner and Schlösser, 2010; Krebs et al., 2011; Boehler et al., 2012; Chiew and Braver, 2013). Consequently, we might expect that working memory would similarly benefit from a reward-induced increase in motivation. A number of studies have examined the influence of monetary incentives on working memory performance. Some of these studies indicate that working memory capacity is enhanced by rewards (Gilbert and Fiez, 2004; Kawasaki and Yamaguchi, 2013), while others find no effect of reward on capacity (Krawczyk et al., 2007; van den Berg et al., 2020). An additional set of studies investigating the relationship between monetary incentives and visual working memory have associated rewards with specific items in the display and have provided more definitive evidence for improved recall precision for rewarded items (Gong and Li, 2014; Klyszejko et al., 2014; Wallis et al., 2015; Thomas et al., 2016; Klink et al., 2017). Under this reward structure, however, monetary incentives are indistinguishable from attentional priority cues. Based on these prior reports, it is unclear how rewards influence the allocation and availability of working memory resources when reward cues are experimentally dissociated from attentional priority cues. That is, do rewards modulate visual working memory performance independent of the prioritization of specific items.

There is reason to believe that working memory performance may not similarly benefit from rewards when incentives are decoupled from attentional priority. Visual working memory is subject to strict capacity limits. Non-rewarded estimates of visual working memory capacity range from 2-4 items (Luck and Vogel, 1997; Cowan, 2001) and capacity estimates are surprisingly stable within an individual (Johnson et al., 2013). Furthermore, capacity limitations have been theorized to arise due to neurophysiological constraints (Lisman and Idiart, 1995; Raffone and Wolters, 2001; Wei et al., 2012; Johnson et al., 2014; Bays, 2015). As such, it may be impossible for visual working memory capacity to be augmented further by the introduction of rewards.

A number of models have been developed to explain capacity limitations. Slot models characterize visual working memory as a small number of indivisible ‘slots’ in which items can be stored (Luck and Vogel, 1997; Cowan, 2001; Awh et al., 2006a; Rouder et al., 2008; Zhang and Luck, 2008). Resource models, on the other hand, consider visual working memory to be instantiated by the allocation of a continuous resource to presented memoranda (Wilken and Ma, 2004; Bays and Husain, 2008; Bays et al., 2009). A central tenet of resource models is that observers are able to flexibly allocate resources across items, such that items can be remembered with variable precision (van den Berg et al., 2012; Fougnie et al., 2012). Recent modeling work suggests that there may also be flexibility in how much total resource can be allocated to a task at hand (van den Berg and Ma, 2018). The amount of resource invested in a particular task is argued to be governed by a tradeoff between the neural cost of using resources and the expected behavioral cost of errors. This would suggest that participants could possibly invest more total resource when there is more at stake. In support of this idea, a recent study found that participants could remember a greater number of items when they were instructed to remember all items in the display rather than focus on a subset of items (Bengson and Luck, 2016). These findings raise the possibility that rewards could similarly increase the total amount of resources available.

Here, we had participants perform a continuous report visual working memory paradigm with independent reward and priority cues. On each trial, participants were presented with a possible reward ($1, $10 or $20) and cue indicating the stimulus that would most likely be probed (e.g. cue valid 80% of the time) following the delay period. We hypothesize 3 possible outcomes for this experiment. One potential result is that monetary incentives have no effect when dissociated from attentional priority (Figure 1B). This result would indicate that prior observations of a reward effect on working memory recall precision could be attributed to prioritization rather than an independent effect of reward. Alternatively, reward could globally enhance working memory representations independent of attentional priority (Figure 1C). In other words, greater reward could result in better recall regardless of whether an item was validly cued or not. This would be consistent with recent work suggesting that rewards attenuate neural noise for both motor and cognitive tasks, leading to more precise responses (Manohar et al., 2015). This result would also suggest that monetary incentives are capable of expanding working memory capacity, consistent with a ‘resource-rational’ theory of working memory (van den Berg and Ma, 2018). Conversely, working memory could present a unique case due to its limited capacity. If working memory capacity cannot be expanded, then rewards could modulate performance primarily through a reallocation of resources according to the attentional priority of presented items. In this case, more accurate memory for prioritized items would come at the expense of less prioritized items. An interaction between cue validity and reward value would provide support for this scenario (Figure 1D).

**Figure 1.**
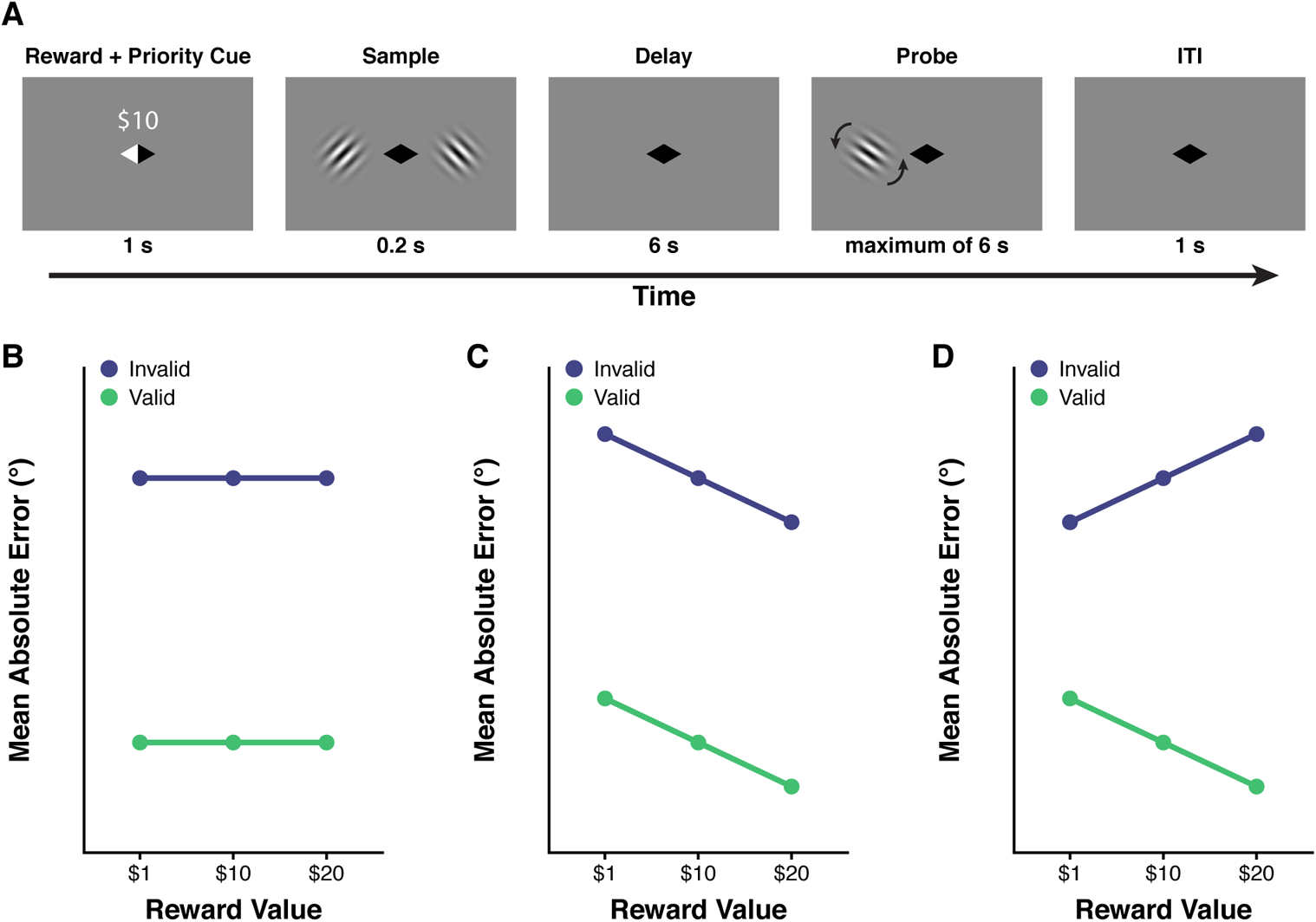
Task paradigm and hypothetical results. (A) Reward and priority continuous report visual working memory paradigm. On each trial, participants were presented with a reward cue ($1, $10 or $20) and a priority cue (left or right hemifield) for 1 s. Participants were then presented with two oriented grating stimuli for 0.2 s. Following a 6 s delay, a probe grating appeared in either the left or right hemifield and participants were given up to 6 s to adjust a probe grating to match the grating that had appeared in the same hemifield. (B-D) Three hypothetical effects of reward and priority. (B) Reward has no effect on visual working memory performance, although there is a robust priority effect across all levels of reward. (C) Reward equally benefits prioritized and de-prioritized items independent of the effect of priority. (D) Reward facilitates performance for prioritized items but this benefit comes at the expense of worse performance for deprioritized items.

## Method

### Participants

92 total healthy adult volunteers participated in this study. 6 participants were unable to finish the experiment due to technical difficulties and were thus excluded from further analysis, leaving us with 86 total participants. All research protocols were approved by the Health Sciences and Behavioral Sciences Institutional Review Board at the University of Michigan. All participants gave written informed consent and were paid $10/hr + performance bonuses for their participation. Participants were recruited from University of Michigan. All participants possessed normal or corrected-to-normal vision. 11 participants participated in Experiment 1 (9 female), 25 participants took part in Experiment 2 (18 female), 25 participants participated in Experiment 3 (17 female), and 25 participants participated in Experiment 4 (18 female).

### Visual Stimuli and Experimental Paradigm

Stimuli were generated and presented using Python with the PsychoPy software package. Participants sat centered in front of a computer screen with a standard QWERTY keyboard. Throughout the session, participants were instructed to fixate on a centrally located black diamond (Figure 1A). Each trial began with the presentation (1 s) of an incentive value ($1, $10, or $20). Participants were instructed that at the end of the experiment, a trial would be selected at random, and if they had accurately reported the probed item (≤ 2° error), they would receive the associated reward for that trial in addition to their base pay. This encouraged participants to evaluate the incentive for each trial independent of all other trials. Simultaneous to the presentation of the reward cue, one side of the fixation diamond changed to white. This cue validly indicated the item that would be probed at the end of the trial on 80% of trials, while on the remaining 20% of trials this cue was invalid. Immediately following the cue period, two circular sine-wave gratings (6 cycles/deg) were presented (one in each hemifield) for 200 ms. The gratings were centered to the left and right of fixation along the horizontal meridian. The orientation of the gratings was drawn from a uniform distribution over 0°-160° in 20° increments with the constraint that the orientation of the gratings could not match. Participants were instructed to then maintain the orientation of the presented gratings over a 6 s delay period during which only the central fixation diamond was visible. Following the delay period, participants were presented with a probe grating that appeared in either the left or right hemifield. The location (hemifield) of the probe grating indicated which sample grating should be recalled. For example, if the probe grating appeared in the left hemifield then participants should report the orientation of the grating that was presented in the left hemifield during the sample display. Participants used the right and left arrow keys to rotate the probe grating clockwise and counter-clockwise, respectively, to match the remembered orientation. Participants were instructed to press the up arrow key to lock in their response and end the current trial. Participants had up to 6 seconds to provide a response. The initial orientation of the probe grating was randomized. The randomization of the probe stimulus ensured that participants were unable to prospectively plan a specific motor action (i.e. direction of rotation) during the maintenance period. Participants were not provided with any feedback at the end of each trial. Trials were separated by a 1 s inter-trial interval.

Subsequent experiments manipulated one of two variables: cue validity or the timing of the priority cue. Experiment 2 served as a direct replication of Experiment 1 and was pre-registered on OSF (https://osf.io/dcjnx). The only differences between the two experiments were that the hemifield cue was valid on 2/3 trials and invalid on the remaining 1/3 trials and the delay period was shortened to 4 s. The change in the level of cue validity enables us to obtain a more reliable estimate of response error on invalid trials (~25 invalid trials per reward value vs. ~12 invalid trials per reward value), with the understanding that the validity effect could potentially be reduced by this change. Experiment 3 examined the effect of reward when the priority cue was completely uninformative (50% valid). Experiment 4 was designed to allow us to isolate the particular phases of working memory where reward modulation might occur. Working memory research has shown that the working memory storage process comprises several phases. These include the initial selection of items, encoding/consolidation of those items into working memory, attentional prioritization of items within working memory (if required), and the maintenance or retention of information over extended delays (Woodman and Vogel, 2004; Awh et al., 2006b; Todd et al., 2011; Myers et al., 2017; Ye et al., 2017). In this experiment, we presented the priority cue 200 ms following the offset of the sample period. If we do not observe an effect of reward, it would suggest that rewards modulate either the initial selection or encoding of information into working memory. In contrast, if we observe the same effect as presenting the priority cue before the sample period, it would suggest that reward modulates processes occurring during either the prioritization of items within working memory or during the maintenance phase.

### Statistical Analysis

All statistical analysis was performed using R (R Core Team, 2019; Version 3.6.1) and MATLAB (Mathworks; Version R2019a). We extracted trial-wise measures of absolute response error for each participant. To model the effect of incentives and cue validity on visual working memory response error we fit a Bayesian hierarchical generalized linear model (GLM). Bayesian statistics provide numerous advantages including allowing one to make intuitive probabilistic statements about the range of credible values (e.g. parameter *x* has a probability of 0.95 of being in the range 1 to 5), as well as incorporate prior knowledge. As absolute response error is strictly positive with positive skew, we specified a Gamma likelihood function and a log link function. The model included fixed effects of reward value ($1, $10, or $20), cue validity (valid or invalid), and their interaction. The model additionally allowed intercepts to vary by subject. A posterior distribution over possible parameter values was sampled using Markov chain Monte Carlo (MCMC) sampling implemented in rstan (Stan Development Team, 2020; Version 2.21.2) via the brms package (Bürkner, 2017, 2018); Version 2.14.4). The model was specified as follows:

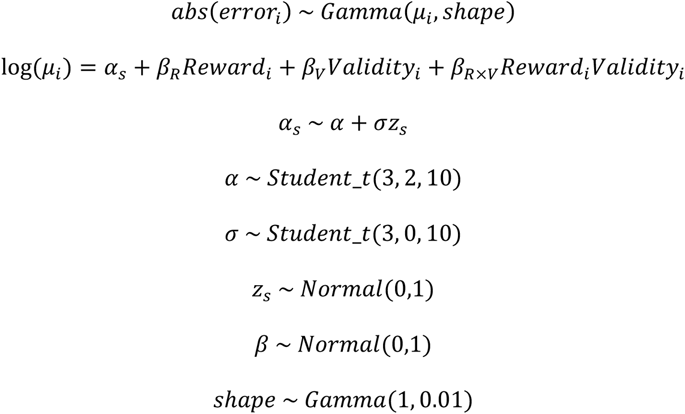

Where *α_s_* denotes a subject-specific intercept and *β_R_, β_V_*, and *β_R×V_* denote parameter estimates for the effect of reward, cue validity, and their interaction. We specified a weakly informative prior distribution for the fixed effects (*β*) with a mean of 0 and a standard deviation of 1 (i.e. N(0,1)). As regression coefficients for a Gamma likelihood GLM with a log link function are interpreted as multiplicative factors rather than slopes, a N(0,1) prior indicates that *a priori* we believe that a change in factor level (e.g. invalid cue to valid cue) is associated with an increase or decrease in absolute error by a factor between 1 and 7.1 (i.e. [exp(0), exp(1.96)]). To ensure that the reported results were robust to the choice of prior, we additionally re-fit the model with both a more liberal (N(0, 4)) and more conservative (N(0, 0.25)) prior for the fixed effects. The overall pattern of effects was robust to the choice of prior. The brms package implements a non-centered parameterization for random effects (Betancourt and Girolami, 2015), which for our model parameterizes subject-specific intercepts (*α_s_*) using an overall intercept (*α*), a subject-specific offset (*Z_s_*), and a scaling parameter (*σ*). This parameterization decorrelates the sampling of random effects from high-order hyperparameters allowing for improved sampling efficiency (Betancourt and Girolami, 2015). Default prior specifications from the brms package were used for parameters associated with subject-specific intercepts (*α*, *Z_s_*, and *σ*), as well as the residual shape parameter (*shape*). Incentive value ($1, $10, or $20) was treated as an ordinal factor.

We ran 4 separate chains with 6000 iterations each. The first 2000 iterations were discarded as warm-up. R-hat values were all very close to 1 (R-hat ≤ 1.001) and effective sample size exceeded 5000 for all parameters indicating that MCMC chains had converged and there was minimal autocorrelation in the sampling. Posterior predictive checks confirmed that distributional assumptions were met and that the specified model could generate data that resembled the actual data. For each parameter in the model, we report the median, 95% highest density interval (HDI), and the probability of direction (*pd*). The HDI represents the interval for which all values within that interval have a higher probability density than points outside that interval. Due to the log-link function, we exponentiate the median and 95% HDI values for reporting so that values represent multiplicative factors on the original response scale (e.g. a one unit change in *x* is associated with an increase in absolute response error by a factor of exp(0.1) = 1.11). *pd* is an index of effect existence (ranging from 50% to 100%), which represents the probability that an effect goes in a particular direction (e.g. effect *x* has a 99% probability of being negative). Note that *pd* represents the probability that the effect is negative or positive prior to exponentiation and the probability that the effect is less than or greater than 1 after exponentiation. We consider a *pd* that is greater than 95% to be strong evidence for the existence of an effect, a *pd* between 80% and 95% to provide some evidence for an effect, and a *pd* that is less than 80% to indicate limited evidence for an effect.

Experiment 2 served as pre-registered replication of the Experiment 1 (https://osf.io/dcjnx). As a result, we performed a Bayesian replication analysis (Ly et al., 2019). This analysis required the pooling of data from both experiments and fitting the same Gamma GLM model. From this analysis we computed a replication Bayes factor, which reflects the evidence for or against a successful replication of the observed effect in the first experiment. To do so, we computed an evidence ratio for both Experiment 1 and the data pooled from both Experiments 1 and 2. The evidence ratio (or Bayes factor) is the ratio of the probability that an effect goes in one direction (e.g. negative) over the probability that it goes in the other direction (e.g. positive) and can be derived using the formula *pd*/(1-*pd*). The evidence ratio we report should not be confused with the Jeffreys Bayes factor (Jeffreys, 1961) that is produced by the R BayesFactor package (Morey & Rouder, 2018) and the statistical software JASP ((Love et al., 2019). The Jeffreys Bayes factor is highly dependent on the definition of the prior. For linear models (*t*-test, linear regression, ANOVA) the range of plausible effect sizes is well known. As a result, default priors can be defined that do not exert undue influence on the resulting Bayes factor (Rouder et al., 2009, 2012; Rouder and Morey, 2012). In other words, the computed value is robust to the particular prior definition used. The choice of prior is not so straightforward for nonlinear models such as those we use here (Rouder et al., 2017). We found that while the HDI for a particular effect was robust to the selected prior definition, the Jeffreys Bayes factor was highly dependent on the prior, ranging from strong evidence for the null to strong evidence for an effect. The replication Bayes factor was computed by dividing the evidence ratio from the pooled model fit by the evidence ratio for Experiment 1 (Ly et al., 2019).

To isolate the specific components of working memory that are modulated by rewards and attentional priority, we additionally performed a mixture model analysis of participants’ response error distribution using the MemToolBox MATLAB package (Suchow et al., 2013). Due to the cue validity manipulation, participants were necessarily presented with far fewer invalid trials. To enable us to properly fit a mixture model to the response error distribution for each condition we collated data from all subjects into a single “super-subject” for each level of reward and cue validity. We fit the following two models: a standard mixture model (Zhang and Luck, 2008) and a swap model (Bays et al., 2009). The standard mixture model fits participants’ response errors with a mixture of two distributions: (1) a uniform distribution reflecting trials in which items were not stored in working memory and (2) a von Mises distribution reflecting the precision of items successfully represented in working memory. This model comprises 2 mixture parameters (which sum to 1), the probability of target report (*P(Target)*) and the probability of guessing (*P(Guess)*), as well as a precision parameter (*SD*). The swap model is an extension of the standard mixture model that includes an additional mixture parameter representing the probability that participants incorrectly reported the identity of the non-cued item [parameters: *P(Target)*, *P(Guess)*, *P(Swap)*, *SD*]. We assessed the relative goodness of fit for each condition using the corrected Akaike Information Criterion (AIC_c_; (Akaike, 1974; Hurvich and Tsai, 1989). For invalid trials of all reward levels in experiment 1, we found lower AIC_c_ values for the swap model relative to the standard mixture model (standard mixture model – swap model AIC_c_ $1, $10, $20: 23.66, 36.10, 22.90), indicating ‘essentially no support’ for the standard mixture model (Burnham and Anderson, 2004). For valid trials, we found support for both models (standard mixture model – swap model AIC_c_: −1.72, −1.53, 4.09). For experiments 2-4, we found no strong preference for one model over the other for both valid and invalid trials (standard mixture model – swap model AIC_c_ range: −2.02 – 5.16). Given the strong preference for the swap model for the Experiment 1 invalid conditions, as well as the fact that the swap model reduces to a standard mixture model when *P(Swap)* is 0, we report results for the swap model for all experiments.

MemToolBox performs an adaptive MCMC sampling procedure (Andrieu et al., 2003), which automatically detects convergence using the (Gelman and Rubin, 1992) statistic. 6000 total samples were collected post-convergence across 3 chains. We computed pairwise comparisons of the posterior distributions for each parameter to assess the effect of priority, reward, and their interaction. To examine the effect of attentional priority we pooled the posterior samples across all levels of reward for invalid and valid trials and then computed all possible pairwise comparisons between the two pooled posterior distributions (324,000,000 comparisons). To test for an interaction between priority and reward, we examined how the difference between valid and invalid posterior parameter distributions changed as a function of reward. To do so, we first computed all pairwise comparisons between posterior samples for invalid and valid trials separately for each level of reward (36,000,000 comparisons each). We then compared the difference distribution for $10 and $20 to the difference distribution for $1. So as to not exceed memory limits by computing all possible pairwise comparisons (1.296 × 10^15^ comparisons), we computed the difference for a random sample of 36,000,000 differences (permute posterior difference distribution for each reward value then subtract one from the other). We then examined whether there was evidence for a tradeoff in the benefit or cost of rewards on prioritized or deprioritized items, respectively. To do so, we computed all possible pairwise comparisons of $10 versus $1 and $20 versus $1 for invalid and valid conditions separately. After flipping the sign of the invalid comparisons, we computed a random sample of 36,000,000 comparisons between the two distributions. As for the hierarchical GLM analysis of absolute error, we report we report the median, 95% HDI, and *pd* for each of these comparisons.

## Results

### Experiment 1

To investigate the effect of monetary incentives on visual working memory, we had participants perform a delayed recall task with oriented contrast gratings (Figure 1A). On each trial, we independently presented a reward cue ($1, $10, or $20) and a stimulus cue (left or right hemifield) prior to stimulus presentation. Participants were told that a trial would be randomly selected at the end of the experiment. If they had accurately reported the probed item (≤ 2° error), they would receive the associated reward for that trial. Critically, rewards were not linked with specific items in the display. In Experiment 1, the stimulus cue validly indicated the to-be-probed item on 80% of trials and invalidly indicated the to-be-probed item on the remaining 20% of trials. The independent manipulation of incentives and cue validity enabled us to examine how reward modulates working memory for stimuli with different levels of attentional priority. We fit a Bayesian hierarchical gamma GLM to trial-wise absolute error with fixed effects of reward value, cue validity, and their interaction.

What is the impact of independent priority and incentive cues on visual working memory recall? Replicating prior work, we found a robust effect of cue validity (Figure 2A; median [95% HDI] = 0.463 [0.400, 0.531], *pd* = 100%), indicating that prioritized items were remembered with greater precision. We further found that the effect of reward depended on whether the priority cue was valid or invalid (Figure 2B; reward × cue validity: median [HDI] = 0.770 [0.611, 0.978], *pd* = 98.5%). Greater incentives were associated with worse recall for invalidly cued trials and better recall for validly cued trials. This interaction suggests that rewards are capable of modulating visual working memory performance. Furthermore, these results indicate that this modulation can be characterized as a reallocation of resources among to-be-remembered items based on their attentional priority (Figure 1D) rather than an increase in the total amount of resources available (Figure 1C).

**Figure 2.**
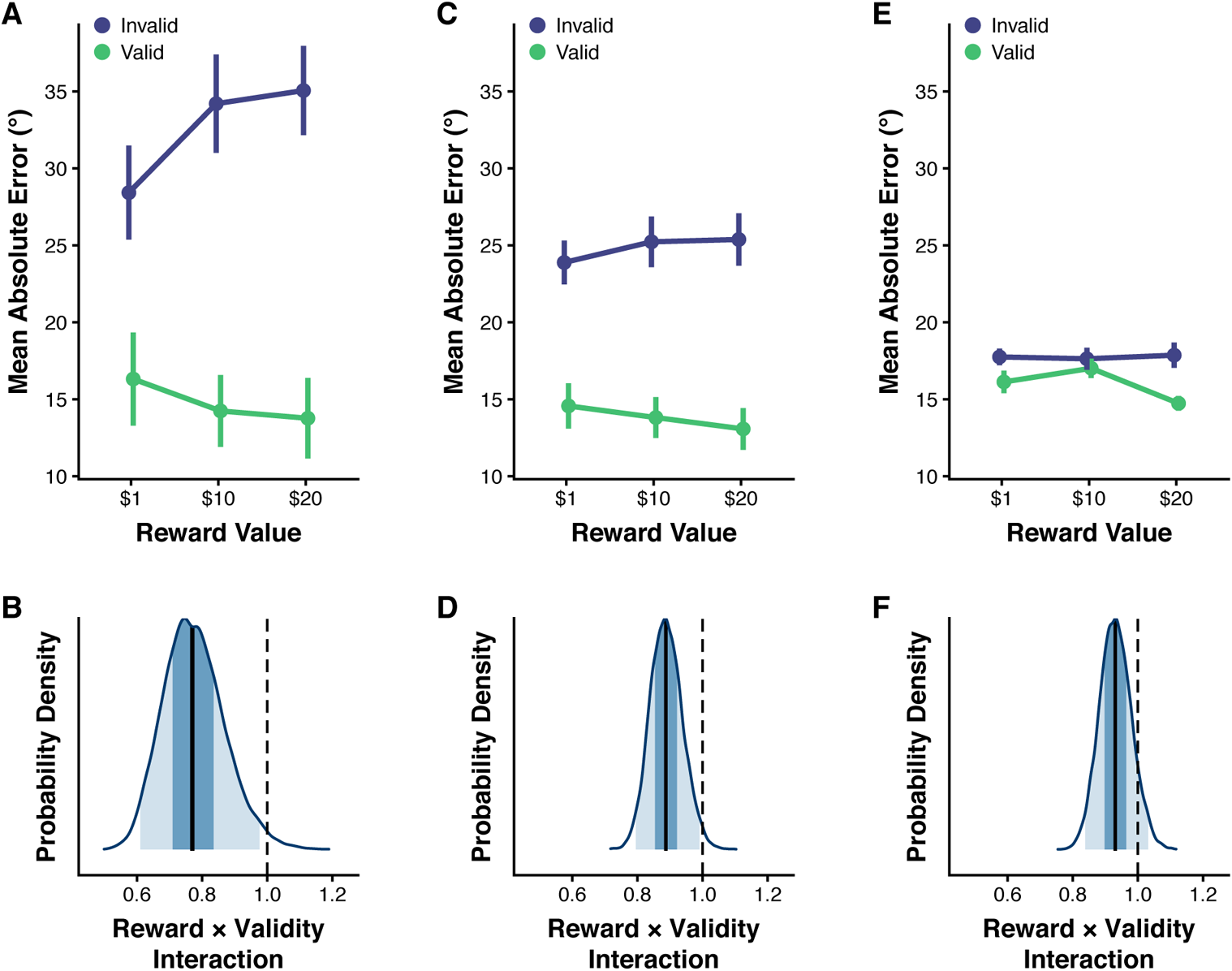
Experiments 1-3 recall error results. (A) Mean absolute recall error across participants for each level of cue validity (valid or invalid) and reward ($1, $10, or $20) in Experiment 1. Error bars represent within-subject standard error of the mean (SEM)(Morey, 2008). (B) Posterior distribution for the interaction between reward and priority for Experiment 1. Posterior parameter values are exponentiated so that they reflect multiplicative factors (with dotted line at 1 representing no change). The dark blue shaded region represents the 50% HDI and the light blue shaded region represents the 95% HDI. (C) Experiment 2 mean absolute recall error for each level of cue validity and reward (D) Experiment 2 posterior distribution for reward × priority interaction. (E) Experiment 3 mean absolute recall error for each level of cue validity and reward. (F) Experiment 3 posterior distribution for reward × priority interaction.

This effect could not be attributed to a speed-accuracy tradeoff. In fact, in addition to worse recall, there was evidence for a slowing of reaction time with increased reward for invalid trials (median [HDI] = 1.096 [1.002, 1.197], *pd* = 97.8%). We found limited evidence for reward modulation of reaction time for valid trials (median [HDI] = 1.019 [0.976, 1.066], *pd* = 79.6%). We did find evidence for a validity effect where participants responded faster on valid trials than invalid trials (median [HDI] = 0.771 [0.728, 0.816], *pd* = 100%).

We next examined whether we could isolate whether the interaction between reward and validity could be tied to a specific component of visual working memory. To do so, we fit a mixture model to a response error distribution aggregated across all participants for each condition. This model comprised 3 mixture parameters (which sum to 1) representing the probability of recalling a target item (*P(Target)*), recalling a non-target item (*P(Swap)*), or guessing randomly (*P(Guess)*) (Bays et al., 2009). The model additionally included an imprecision parameter, *SD*, which represents the standard deviation of recall error for trials for which the participant successfully remembered an item (target or non-target). We tested for a validity effect by computing all pairwise differences between posterior distributions for valid and invalid trials yielding a posterior probability difference distribution.

We found strong evidence for a validity effect for the probability of target report (*P(Target)*: median [HDI] = 0.416 [0.118, 0.714], *pd* = 99.86%; see Table 1) and the probability of an item swap (*P(Swap)*: median [HDI] = −0.334 [−0.529, −0.167], *pd* = 100%). There was weaker evidence for a validity effect for recall precision (*SD*: median [HDI] = −5.507 [−15.026, 4.512], *pd* = 83.75%) and little evidence for a validity effect for the probability of guessing (*P(Guess)*: median [HDI] = −0.052 [−0.429, 0.251], *pd* = 61.54%).

**Table 1.** Mixture Model Posterior Summary for Experiment 1

| Validity | Parameter | \$1 | | \$10 | | \$20 | |
| --- | --- | --- | --- | --- | --- | --- | --- |
|  |  | Median | 95% HDI | Median | 95% HDI | Median | 95% HDI |
| Valid | $P(\textit{Target})$ | 0.72 | 0.66 – 0.78 | 0.85 | 0.8 – 0.9 | 0.87 | 0.83–0.91 |
| | $P(\textit{Guess})$ | 0.26 | 0.21 – 0.33 | 0.14 | 0.08 – 0.19 | 0.1 | 0.05 – 0.15 |
| | $P(\textit{Swap})$ | 0.01 | 0.00 – 0.04 | 0.01 | 0.00 – 0.04 | 0.03 | 0.00 – 0.06 |
| | $SD$ | 11.7 | 9.74 – 13.08 | 14.53 | 12.71 – 16.23 | 14.41 | 13.18 – 16.08 |
| Invalid | $P(\textit{Target})$ | 0.55 | 0.33 – 0.68 | 0.38 | 0.25 – 0.53 | 0.27 | 0.11 – 0.51 |
| | $P(\textit{Guess})$ | 0.13 | 0.00 – 0.35 | 0.18 | 0.02 – 0.41 | 0.4 | 0.05 – 0.67 |
| | $P(\textit{Swap})$ | 0.31 | 0.18 – 0.45 | 0.42 | 0.27 – 0.56 | 0.34 | 0.18 – 0.54 |
| | $SD$ | 19.89 | 10.86 – 27.11 | 19.53 | 12.19 – 25.87 | 16.5 | 8.91 – 30.32 |

We next tested for an interaction by comparing the difference between valid and invalid trials for different levels of reward. An interaction would be evidenced by a larger difference between levels of cue validity at $10 and $20 than $1. We found that the effect of attentional priority on the probability of recalling the target item or guessing was modulated by reward (Figure 3A-B; *P(Target) [$10 V – I] vs [$1 V – I]*: median [HDI] = 0.281 [0.024, 0.516], *pd* = 98.22%; *P(Target) [$20 V – I] vs [$1 V – I]*: median [HDI] = 0.410 [0.107, 0.673], *pd* = 99.45%; *P(Guess) [$10 V – I] vs [$1 V – I]*: median [HDI] = −0.176 [−0.485, 0.140], *pd* = 86.29%; *P(Guess) [$20 V – I] vs [$1 V – I]*: median [HDI] = −0.408 [−0.759, 0.013], *pd* = 96.37%). We found little evidence for an interaction between priority and reward for the probability of a memory swap or imprecision (Supplementary 1A-B; *P(Swap) [$10 V – I] vs [$1 V – I]*: median [HDI] = −0.104 [−0.312, 0.112], *pd* = 82.69%; *P(Swap) [$20 V – I] vs [$1 V – I]*: median [HDI] = 0.007 [−0.254, 0.220], *pd* = 52.45%; *SD [$10 V – I] vs [$1 V – I]*: median [HDI] = 3.148 [−8.256, 14.403], *pd* = 70.09%; *SD [$20 V – I] vs [$1 V – I]*: median [HDI] = 5.283 [−10.435, 18.392], *pd* = 72.92%). Thus, rewards appear to primarily affect the success of encoding items into working memory rather than the probability of item swaps in working memory or the precision of working memory representations.

**Figure 3.**
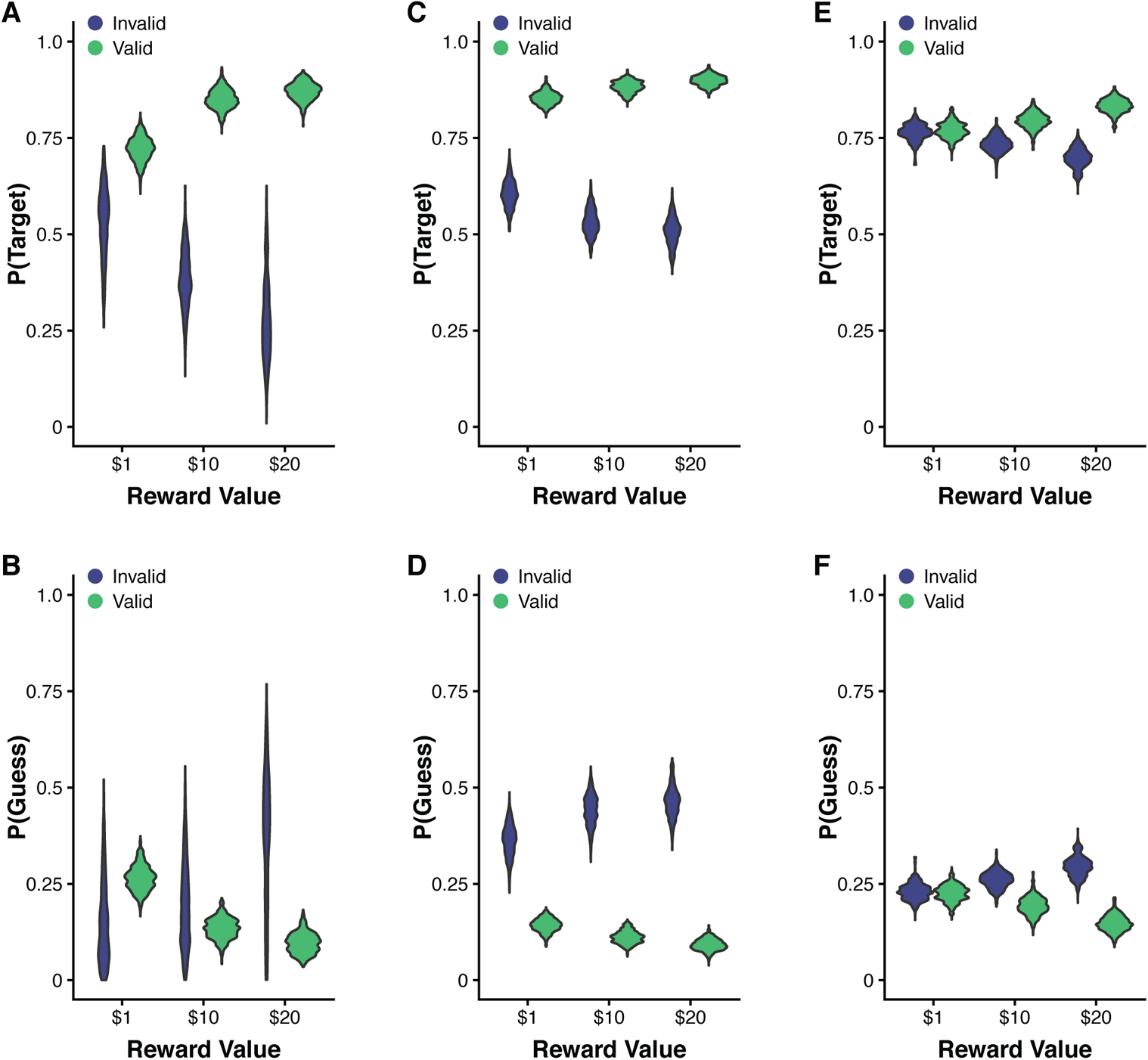
Experiments 1-3 mixture model results. Violin plots depict posterior distributions for mixture model parameters for each level of cue validity and reward. (A) Experiment 1 probability of target report posterior distributions. (B) Experiment 1 probability of guessing posterior distributions. (C) Experiment 2 probability of target report posterior distributions. (D) Experiment 2 probability of guessing posterior distributions. (E) Experiment 3 probability of target report posterior distributions. (F) Experiment 3 probability of guessing posterior distributions.

We then examined whether there was a tradeoff in the probability of target recall or guessing between prioritized and de-prioritized items. In other words, we examine whether any benefit associated with reward for prioritized items was accompanied by a cost for de-prioritized items. We tested for a tradeoff by comparing the difference between larger reward values ($10 and $20) and low reward ($1) for valid and invalid items to see if the absolute difference was of similar magnitude. We found little evidence for an asymmetry in the difference between larger reward values and low reward between validly and invalidly cued trials (*P(Target) [Valid $10 – $1] vs. [Invalid $1 – $10]*: median [HDI] = −0.022 [−0.257, 0.235], *pd* = 56.75%; *P(Target) [Valid $20 – $1] vs. [Invalid $1 – $20]*: median [HDI] = −0.112 [−0.374, 0.191], *pd* = 75.94%; *P(Guess) [Valid $10 – $1] vs. [Invalid $1 – $10]*: median [HDI] = −0.085 [−0.400, 0.225], *pd* = 70.58%; *P(Guess) [Valid $20 – $1] vs. [Invalid $1 – $20]*: median [HDI] = 0.071 [−0.350, 0.423], *pd* = 62.21%). So we again find evidence that rewards do not increase the total amount of encoded information. Rather, incentive cues influence how displayed items are prioritized for subsequent encoding.

### Experiment 2

In Experiment 2, we aimed to replicate Experiment 1 in a larger sample of participants (N = 25) to confirm that reward primarily influences working memory task performance by modulating attentional priority. In order to collect a greater number of invalid trials and obtain a more accurate estimate of response error, cue validity was reduced to 66.67% of trials (versus 80% in Experiment 1) and the delay period was shortened to 4 s (versus 6 s in Experiment 1). The task was otherwise identical to that used in Experiment 1.

Participants were far more accurate (i.e., lower mean absolute error) on valid trials than invalid trials (Figure 2C; median [HDI] = 0.559 [0.524, 0.594], *pd* = 100%), again indicating that the priority cue manipulated attentional priority (Figure 2B). We found strong evidence for an interaction between reward and cue validity (Figure 2D; median [HDI] = 0.887 [0.793, 0.987], *pd* = 98.4%), whereby the memory advantage for validly cued items was larger at the higher reward values. Again, this interaction could not be explained by a speed-accuracy tradeoff, as there was no evidence for an effect of reward on reaction time for both valid and invalid trials (valid: median [HDI] = 1.010 [0.978, 1.041], *pd* = 72.2%; invalid: median [HDI] = 0.991 [0.970, 1.013], *pd* = 78.8%). These results further demonstrate that rewards affect working memory primarily through a change in the degree of prioritization among to-be-remembered items.

We next performed a Bayesian replication analysis to quantify the evidence for or against a successful replication (Ly et al., 2019). To do so, we computed an evidence updating replication Bayes factor, which involves fitting the same model to data pooled from both experiments (N = 36) and then comparing the resulting evidence to the evidence from the original experiment (see methods). The interaction between reward and cue validity for Experiment 1 yielded an evidence ratio of 65.67. The pooled model fit revealed the same interaction between reward and cue validity (median [HDI] = 0.868 [0.787, 0.958], *pd* = 99.8%, evidence ratio = 499). This resulted in a replication Bayes factor of 7.60 (i.e. 499 / 65.67), which indicates that the data from the replication attempt (Experiment 2) are 7.60 times more likely under the hypothesis that the effect is consistent with the one found in the original study (Experiment 1) than the null hypothesis that the effect is not consistent.

We again performed a mixture model analysis to determine which aspect of working memory was modulated by reward and priority (see Table 2). We found strong evidence of a validity effect on the probability of target recall and guessing (*P(Target) V – I*: median [HDI] = 0.334 [0.211, 0.440], *pd* = 100%; *P(Guess) V – I*: median [HDI] = −0.311 [−0.426, −0.177], *pd* = 100%). We found weaker evidence for an effect of cue validity on the probability of a memory swap (*P(Swap) V – I*: median [HDI] = −0.020 [−0.065, 0.010], *pd* = 89.49%) and there was little evidence for an effect of cue validity on recall precision (*SD V – I*: median [HDI] = −0.651 [−4.146, 2.892], *pd* = 63.59%). Replicating Experiment 1, we observed a strong interaction between reward and priority for both the probability of target recall and the probability of guessing (Figure 3C-D; *P(Target) [$10 V – I] vs [$1 V – I]*: median [HDI] = 0.103 [−0.005, 0.207], *pd* = 96.87%; *P(Target) [$20 V – I] vs [$1 V – I]*: median [HDI] = 0.143 [0.036, 0.253], *pd* = 99.59%; *P(Guess) [$10 V – I] vs [$1 V – I]*: median [HDI] = −0.110 [−0.230, 0.011], *pd* = 96.23%; *P(Guess) [$20 V – I] vs [$1 V – I]*: median [HDI] = −0.149 [−0.273, −0.030], *pd* = 99.39%). There was little evidence for an interaction between reward and priority for the probability of non-target report or recall precision (Supplementary Figure 1C-D; *P(Swap) [$10 V – I] vs [$1 V – I]*: median [HDI] = 0.007 [−0.048, 0.065], *pd* = 61.08%; *P(Swap) [$20 V – I] vs [$1 V – I]*: median [HDI] = 0.007 [−0.048, 0.064], *pd* = 59.37%; *SD [$10 V – I] vs [$1 V – I]*: median [HDI] = 1.454 [−2.263, 5.251], *pd* = 77.34%; *SD [$20 V – I] vs [$1 V – I]*: median [HDI] = 2.164 [−1.823, 6.197], *pd* = 86.02%). We further found evidence for a tradeoff in the effect of reward on probability of target recall and guessing between cued and non-cued items. The change in probability associated with larger rewards for valid trials (e.g. increased probability of target report) was mirrored by a change in the opposite direction for invalid trials (e.g. decreased probability of target report) (*P(Target) [V $10 – $1] vs. [I $1 – $10]*: median [HDI] = −0.0447 [−0.150, 0.062], *pd* = 79.12%; *P(Target) [V $20 – $1] vs. [I $1 – $20]*: median [HDI] = −0.052 [−0.162, 0.055], *pd* = 82.90%; *P(Guess) [V $10 – $1] vs. [I $1 – $10]*: median [HDI] = 0.048 [−0.073, 0.168], *pd* = 77.64%; *P(Guess) [V $20 – $1] vs. [I $1 – $20]*: median [HDI] = 0.048 [−0.072, 0.171], *pd* = 78.14%). These findings further indicate that rewards do not change the availability of resources. Instead, rewards appear to modulate the distribution of resources across to-be-remembered items.

**Table 2.** Mixture Model Posterior Summary for Experiment 2

| Validity | Parameter | \$1 | | \$10 | | \$20 | |
| --- | --- | --- | --- | --- | --- | --- | --- |
|  |  | Median | 95% HDI | Median | 95% HDI | Median | 95% HDI |
| Valid | $P(\textit{Target})$ | 0.85 | 0.82 – 0.88 | 0.88 | 0.85 – 0.91 | 0.9 | 0.87 – 0.93 |
| | $P(\textit{Guess})$ | 0.14 | 0.11 – 0.17 | 0.11 | 0.08 – 0.15 | 0.09 | 0.06 – 0.12 |
| | $P(\textit{Swap})$ | 0.002 | 0.00 – 0.01 | 0.005 | 0.00 – 0.02 | 0.008 | 0.00 – 0.02 |
| | $SD$ | 14.82 | 13.88 – 15.69 | 15.12 | 13.93 – 15.94 | 14.33 | 13.48 – 15.24 |
| Invalid | $P(\textit{Target})$ | 0.61 | 0.54 – 0.67 | 0.53 | 0.47 – 0.59 | 0.51 | 0.44 – 0.58 |
| | $P(\textit{Guess})$ | 0.37 | 0.29 – 0.45 | 0.44 | 0.36 – 0.51 | 0.46 | 0.40 – 0.56 |
| | $P(\textit{Swap})$ | 0.03 | 0.00 – 0.07 | 0.02 | 0.00 – 0.06 | 0.03 | 0.00 – 0.06 |
| | $SD$ | 16.57 | 14.50 – 19.20 | 15.43 | 13.13 – 17.84 | 13.95 | 11.11 – 16.86 |

### Experiment 3

In Experiments 1 and 2, the greater allocation of resources to prioritized items could be construed as a strategy to allocate resources to items with higher expected value. Experiment 3 examined the effect of reward on the allocation of working memory resources when the cued item no longer had greater expected value. To do so, we further manipulated cue validity so that only 50% of priority cues were valid. Prioritizing the cued item in this case would not lead to any overall benefits in performance. We again found strong evidence for a validity effect (Figure 2E; median [HDI] = 0.885 [0.836, 0.940], *pd* = 100%) as well as some modulation of this validity effect by reward (Figure 2F; median [HDI] = 0.931 [0.841, 1.033], *pd* = 91.5%). However, we surprisingly did not find strong evidence for a cost. Greater rewards were associated with reduced recall error for valid trials (median [HDI] = 0.919 [0.856, 0.990], *pd* = 98.8%), but there was little evidence for a change in recall error with increasing reward for invalid trials (median [HDI] = 0.987 [0.918, 1.061], *pd* = 64.4%). Nevertheless, we again find evidence for an asymmetry in the influence of reward depending on the attentional priority of the remembered item.

We then fit a mixture model to the response error distribution for each trial type (see Table 3). This analysis allowed us to distinguish between different sources of recall error. We found some evidence for a validity effect for the probability of target report and the probability of guessing (*P(Target) Valid – Invalid*: median [HDI] = 0.066 [−0.027, 0.161], *pd* = 91.57%; *P(Guess) Valid – Invalid*: median [HDI] = −0.072 [−0.176 0.026], *pd* = 91.63%), as well as strong evidence for an interaction between reward and cue validity for *P(Target)* and *P(Guess)* (Figure 3E-F; *P(Target) [$10 V – I] vs [$1 V – I]*: median [HDI] = 0.055 [−0.031, 0.138], *pd* = 89.60%; *P(Target) [$20 V – I] vs [$1 V – I]*: median [HDI] [HDI] = 0.130 [0.043, 0.216], *pd* = 99.73%; *P(Guess) [$10 V – I] vs [$1 V – I]*: median [HDI] = −0.064 [−0.150, 0.026], *pd* = 91.68%; *P(Guess) [$20 V – I] vs [$1 V – I]*: median [HDI] = −0.143 [−0.234, −0.051], *pd* = 99.82%). There was limited evidence for a validity effect or an interaction between reward and validity for the probability of a memory swap (Supplementary Figure 1E; all *pd* < 83%). There was evidence for an interaction between reward and validity for recall precision (Supplementary Figure 1F; *pd* > 96.5%). However, the direction of this effect was opposite to that observed for the probability of target report and guessing (i.e. larger difference between valid and invalid trials for $1 than $10 or $20). In contrast to the analysis of absolute error, we did find evidence for a tradeoff whereby the benefit observed with increasing reward for the valid cue condition was accompanied by a cost in the opposite direction for the invalid cue condition. This tradeoff was observed for both the probability of target recall and the probability of guessing (*P(Target) [Valid $10 – $1] vs. [Invalid $1 – $10]*: median [HDI] = −0.005 [−0.089, 0.080], *pd* = 54.99%; *P(Target) [Valid $20 – $1] vs. [Invalid $1 – $20]*: median [HDI] = −0.002 [−0.089, 0.084], *pd* = 52.08%; *P(Guess) [Valid $10 – $1] vs. [Invalid $1 – $10]*: median [HDI] = −0.002 [−0.091, 0.085], *pd* = 52.17%; *P(Guess) [Valid $20 – $1] vs. [Invalid $1 – $20]*: median [HDI] = −0.014 [−0.105, 0.077], *pd* = 62.08%). These findings suggest that reward modulation of prioritization may be independent of any strategic allocation of resources to items with greater expected value.

**Table 3.** Mixture Model Posterior Summary for Experiment 3

| Validity | Parameter | \$1 | | \$10 | | \$20 | |
| --- | --- | --- | --- | --- | --- | --- | --- |
|  |  | Median | 95% HDI | Median | 95% HDI | Median | 95% HDI |
| Valid | $P(\textit{Target})$ | 0.77 | 0.72 – 0.81 | 0.79 | 0.75 – 0.83 | 0.83 | 0.79 – 0.87 |
| | $P(\textit{Guess})$ | 0.23 | 0.18 – 0.27 | 0.19 | 0.14 – 0.23 | 0.15 | 0.11 – 0.19 |
| | $P(\textit{Swap})$ | 0.003 | 0.00 – 0.01 | 0.01 | 0.00 – 0.03 | 0.02 | 0.00 – 0.04 |
| | $SD$ | 13.58 | 12.22 – 14.66 | 14.71 | 13.65 – 15.95 | 13.13 | 12.08 – 14.49 |
| Invalid | $P(\textit{Target})$ | 0.76 | 0.72 – 0.80 | 0.73 | 0.70 – 0.78 | 0.7 | 0.65 – 0.74 |
| | $P(\textit{Guess})$ | 0.23 | 0.19 – 0.27 | 0.26 | 0.22 – 0.30 | 0.29 | 0.25 – 0.35 |
| | $P(\textit{Swap})$ | 0.004 | 0.00 – 0.02 | 0.004 | 0.00 – 0.02 | 0.006 | 0.00 – 0.02 |
| | $SD$ | 15.00 | 13.54 – 16.55 | 13.82 | 12.32 – 15.02 | 12.13 | 11.14 – 13.31 |

### Experiment 4

We next sought to isolate the specific phase of working memory that is modulated by rewards. To do so, rather than present the stimulus cue concurrently with the reward cue, we shifted the stimulus cue to appear immediately following the offset of the stimulus presentation period. This post-cue could only affect prioritization after encoding had already occurred. This change allowed us to determine if rewards modulate processing during selection/encoding phases of the trial or maintenance/retrieval phases of the trial.

We again found evidence for a validity effect (median [HDI] = 0.903 [0.849, 0.961], *pd* = 100%). In contrast to Experiments 1-3, we found limited evidence for an interaction between reward and validity (Figure 4A-B; median [HDI] = 0.974 [0.877, 1.081], *pd* = 69.5%). The lack of an interaction indicates that rewards do not modulate performance once items have already been encoded into working memory. This would suggest that the reward modulation observed in Experiments 1-3 must occur during either the attentional selection of items or the subsequent encoding of those items into working memory. This finding was confirmed by a mixture model analysis (see Table 4). While we found some evidence for an effect of cue validity on the probability of target recall (*P(Target) Valid – Invalid*: median [HDI] = 0.065 [−0.035, 0.153], *pd* = 88.92%), we did not find strong evidence for an interaction between reward and cue validity for any model parameter when the priority cue was retroactive (Figure 4C-D; Supplementary Figure 2A-B; all *pd* < 90%). These results again indicate that rewards do not influence working memory performance when participants are unable to prioritize information prior to encoding.

**Figure 4.**
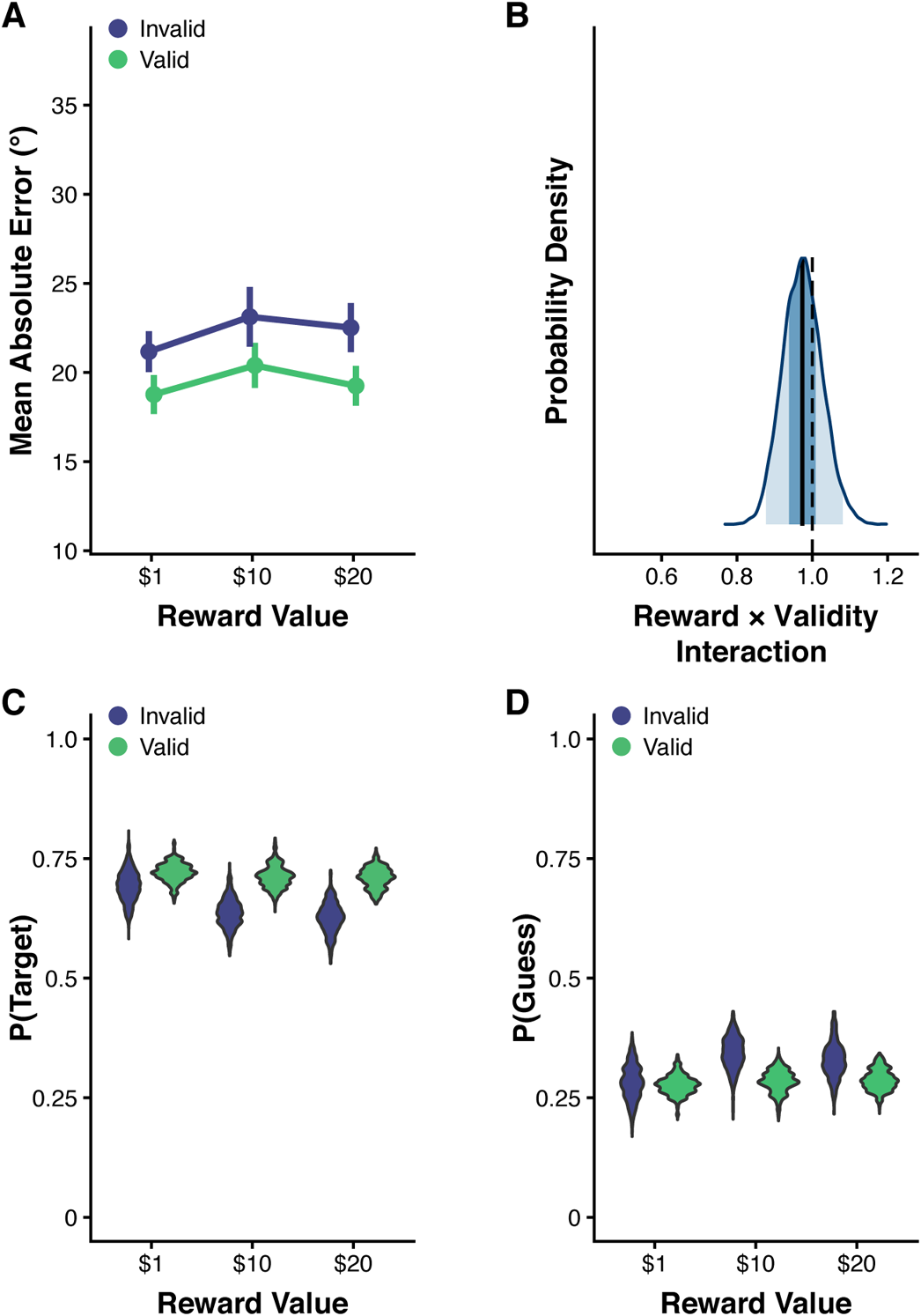
Experiment 4 results. (A) Mean absolute recall error across participants for each level of cue validity and reward. Error bars represent within-subject SEM. (B) Posterior distribution for the interaction between reward and priority. The dark blue shaded region represents the 50% HDI and the light blue shaded region represents the 95% HDI. (C) Violin plot depicting the posterior distribution for the probability of target report for each level of cue validity and reward. (D) Posterior distributions for the probability of guessing.

**Table 4.** Mixture Model Posterior Summary for Experiment 4

| Validity | Parameter | \$1 | | \$10 | | \$20 | |
| --- | --- | --- | --- | --- | --- | --- | --- |
|  |  | Median | 95% HDI | Median | 95% HDI | Median | 95% HDI |
| Valid | $P(\textit{Target})$ | 0.72 | 0.68 – 0.75 | 0.71 | 0.67 – 0.76 | 0.71 | 0.67 – 0.75 |
| | $P(\textit{Guess})$ | 0.28 | 0.24 – 0.32 | 0.28 | 0.24 – 0.33 | 0.29 | 0.25 – 0.34 |
| | $P(\textit{Swap})$ | 0.001 | 0.00 – 0.01 | 0.003 | 0.00 – 0.01 | 0.002 | 0.00 – 0.01 |
| | $SD$ | 15.10 | 14.03 – 16.26 | 16.47 | 15.14 – 17.98 | 14.80 | 13.73 – 15.95 |
| Invalid | $P(\textit{Target})$ | 0.69 | 0.63 – 0.75 | 0.64 | 0.58 – 0.70 | 0.63 | 0.57 – 0.69 |
| | $P(\textit{Guess})$ | 0.28 | 0.22 – 0.36 | 0.34 | 0.28 – 0.41 | 0.33 | 0.26 – 0.41 |
| | $P(\textit{Swap})$ | 0.02 | 0.00 – 0.06 | 0.02 | 0.00 – 0.05 | 0.04 | 0.01 – 0.09 |
| | $SD$ | 16.00 | 13.86 – 18.50 | 15.60 | 13.81 – 17.33 | 14.99 | 13.33 – 17.42 |

## Discussion

The effect of motivation on visual working memory is controversial. Working memory is subject to strict capacity limits that may restrict the potential benefits of reward on performance (Luck and Vogel, 1997; Cowan, 2001). On the other hand, recent modeling work theorizes that the total amount of resource dedicated to storing items in working memory is flexible and potentially able to be increased when there is more at stake (van den Berg and Ma, 2018). Prior studies on the effect of motivation on visual working memory capacity and precision have produced mixed results (Gilbert and Fiez, 2004; Krawczyk et al., 2007; Kawasaki and Yamaguchi, 2013; van den Berg et al., 2020). Here, we find that rewards do modulate working memory performance. However, the direction and extent of this modulation was determined by the attentional priority of the to-be-remembered item. Our findings provide evidence for flexibility in resource allocation in the selection of information for working memory. Resource availability, however, is unaffected by the prospect of reward.

We independently manipulated reward and attentional priority in a visual working memory task and found that greater rewards coincided with greater prioritization of pre-cued items at the expense of non-cued items. This was evidenced by a decrease in recall error for cued items and an increase in recall error for non-cued items with larger rewards. We additionally performed a mixture model analysis that enabled us to disentangle the effects of reward on different components of visual working memory. We found that rewards modulated the probability of successfully encoding an item into working memory. Greater rewards were associated with greater probability of successful encoding for cued items and decreased probability of successful encoding for non-cued items. Interestingly, we found little evidence that rewards modulated the probability of binding errors or recall precision. This is consistent with a recent study that found no effect of reward on visual working memory precision (van den Berg et al., 2020). Taken together, these findings indicate that monetary incentives primarily affect whether an item is prioritized for encoding rather than the quality of visual working memory representations or encoding errors (e.g. misbinding of location and feature).

Our results provide support for a proactive control account of reward modulation of visual working memory. Proactive control refers to the advance maintenance of goal-relevant information for the purpose of biasing perception-, attention-, and action-related systems (Braver, 2012). Proactive control can be differentiated from reactive control which is characterized by the transient adjustment of control in response to unexpected events (Braver, 2012). Prior work has indicated that incentives primarily enhance proactive rather than reactive control processes (Braver et al., 2009; Chiew and Braver, 2013, 2014, 2016). For example, Chiew and Braver (2016) independently manipulated incentives and task-relevant cues and found that rewards modulate performance specifically when preparatory cues are presented, but only when there is sufficient time to act on those cues. This is consistent with the current result that incentives modulate the priority of displayed items, but only when priority cues are presented prior to the encoding period. Once participants had encoded items into visual working memory (Experiment 4), prospective rewards no longer impacted performance. This is notable as it indicates that rewards do not modulate prioritization during maintenance. These findings potentially suggest distinct mechanisms may underlie prioritization of sensory stimuli versus the prioritization of items held in working memory.

Our results cannot be explained by a strategy whereby participants simply prioritize items with greater expected value (Shenhav et al., 2013, 2016). In Experiments 1 and 2, it could be argued that cued and non-cued items had unequal expected value (e.g. 0.8 * $20 = $16 *vs.* 0.2 * $20 = $4). As a result, participants could simply have strategically deployed greater attentional resources to the item with greater value. In Experiment 3, however, priority cues had equal chance of being valid or invalid. Thus, neither displayed item had greater expected value. Nevertheless, we still observed an interaction between reward and priority such that greater incentives were associated with greater likelihood that the cued item would be encoded and decreased likelihood that the non-cued item would be encoded. This finding suggests that reward modulation of prioritization may be an involuntary process akin to the phenomenon of value-driven attentional capture (Anderson et al., 2011). Recent conceptualizations of the attentional capture effect suggest that attention can be captured by salient stimuli but the degree of capture can be modulated or even prevented if the attentional control system is appropriately configured (Luck et al., 2021). As a result, both salience and control settings influence the priority of specific locations and features. The current findings potentially suggest that rewards produce an involuntary reconfiguration of control that increases the degree to which cues drive resource allocation regardless of whether they are informative or not.

Our results argue against a resource rational model of visual working memory (van den Berg and Ma, 2018). According to the resource rational model, an observer could potentially expend greater neural resources when there is greater value at stake, and thus increase storage capacity. We found little evidence for a change in the availability of resources with greater rewards. Instead, rewards simply modulated the flexible allocation of resources to different items within the display. The current results indicate that the effect of reward on visual working memory differs from the primarily facilitatory effect previously shown for motor processes or higher order decision processes (Takikawa et al., 2002; Hübner and Schlösser, 2010; Krebs et al., 2011; Boehler et al., 2012; Chiew and Braver, 2013, 2016; Manohar et al., 2015, 2018; Anderson et al., 2020; Adkins et al., 2021). Rather than providing a global boost to performance, any reward benefit for prioritized items came at the expense of deprioritized items. While the prospect of reward influences proactive control at encoding, visual working memory capacity constraints appear to be unaffected.

## Acknowledgements

We thank Katy Michon for help with data collection. This work was funded by a National Institutes of Health grant (F32MH124268 to J.A.B.). The views expressed in this article do not necessarily represent the views of the NIH or the United States Government. The authors declare no competing financial interests

---

**Supplementary Figure 1.**
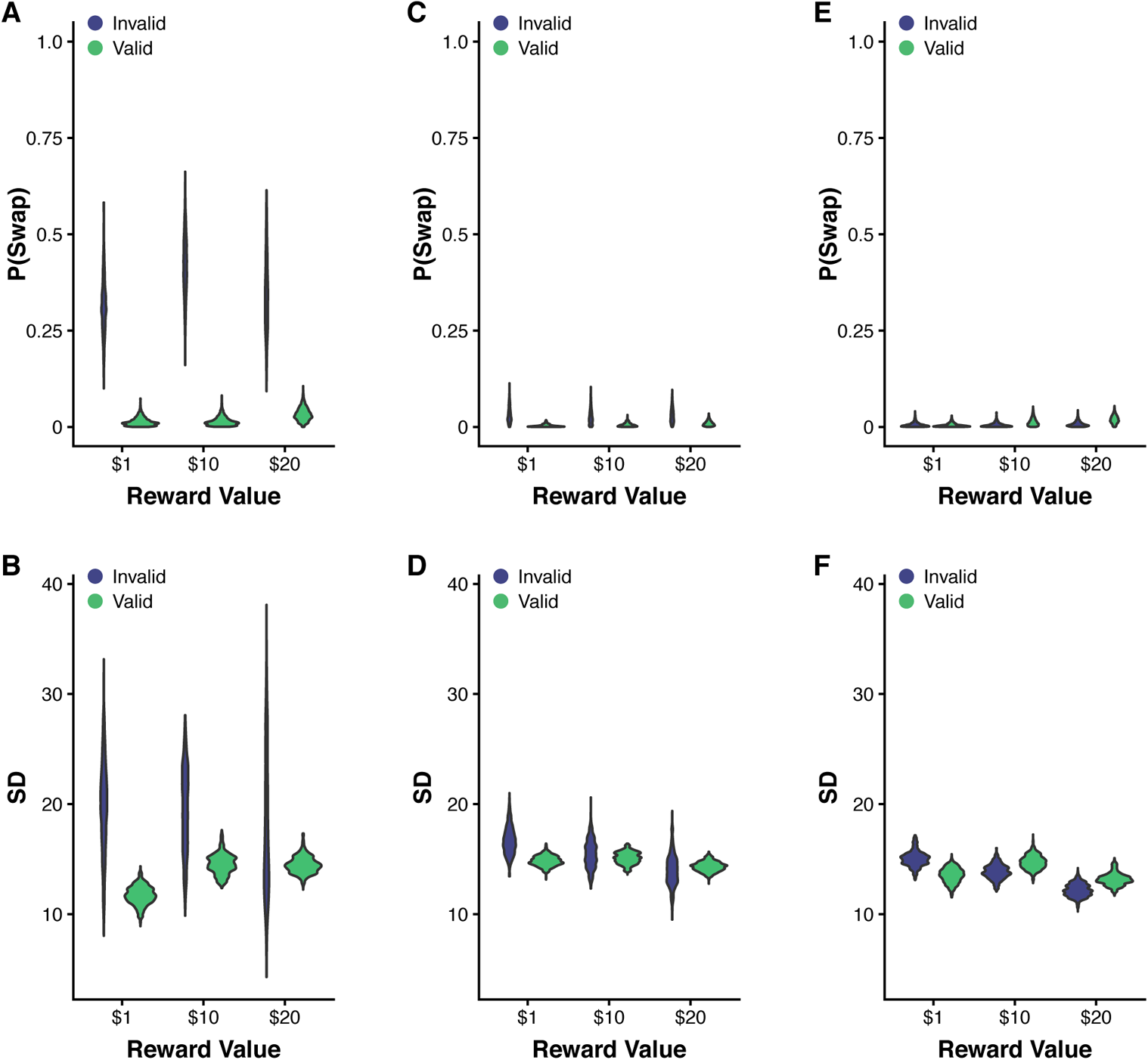
Experiments 1-3 mixture model results. Violin plots depict posterior distributions for mixture model parameters for each level of cue validity and reward. (A) Experiment 1 probability of memory swap posterior distributions. (B) Experiment 1 recall imprecision posterior distributions. (C) Experiment 2 probability of memory swap posterior distributions. (D) Experiment 2 recall imprecision posterior distributions. (E) Experiment 3 probability of memory swap posterior distributions. (F) Experiment 3 recall imprecision posterior distributions.

**Supplementary Figure 2.**
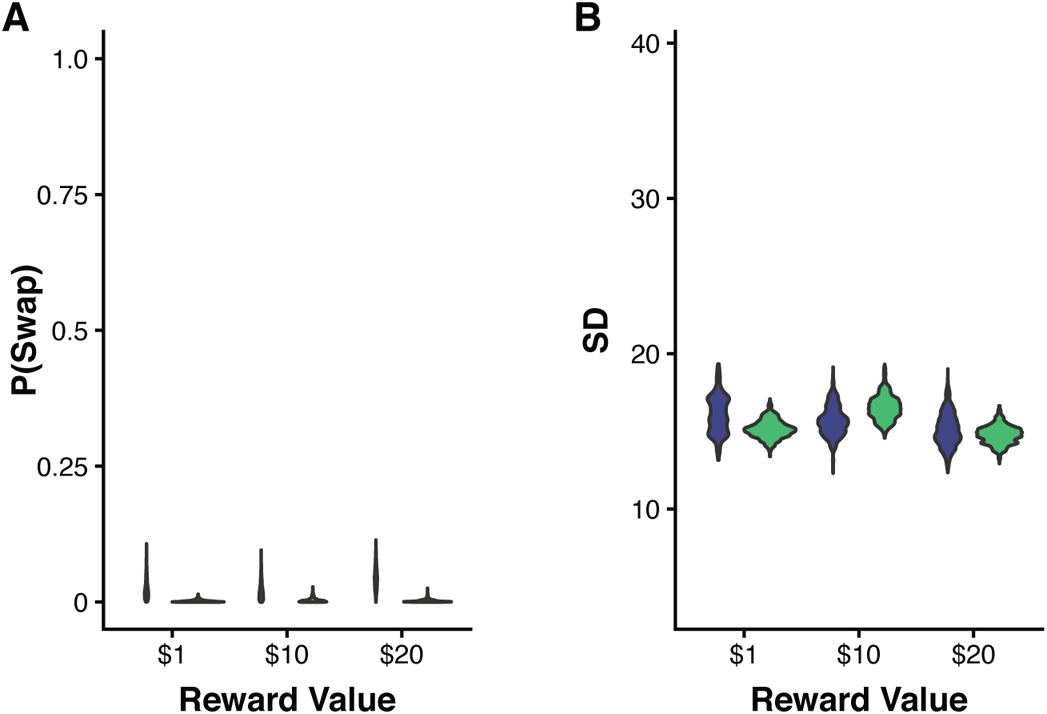
Experiment 4 mixture model results. Violin plots depict posterior distributions for mixture model parameters for each level of cue validity and reward. (A) Probability of memory swap posterior distributions. (B) Recall imprecision posterior distributions.

